# Single extracellular vesicle analysis in human amniotic fluid shows evidence of phenotype alterations in preeclampsia

**DOI:** 10.1101/2022.02.14.480331

**Authors:** Natalia Gebara, Julia Scheel, Renata Skovronova, Cristina Grange, Luca Marozio, Shailendra Gupta, Veronica Giorgione, Federico Caicci, Chiara Benedetto, Asma Khalil, Benedetta Bussolati

**Affiliations:** Department of Molecular Biotechnology and Health Sciences, University of Turin, Italy; Department of Systems Biology and Bioinformatics, University of Rostock, Germany; Department of Medical Sciences, University of Turin, Turin, Italy; Department of Surgical Sciences, Obstetrics and Gynecology, University of Turin, Italy; Vascular Biology Research Centre, Molecular and Clinical Sciences Research Institute, St George’s University of London, UK; DiBio Imaging facility, University of Padua, Italy; Fetal Medicine Unit, St George’s University Hospitals NHS Foundation Trust, St George’s University of London, UK

**Keywords:** extracellular vesicles, exosomes, amniotic fluid, placenta, pregnancy disorders, angiogenesis, soluble endoglin

## Abstract

Amniotic fluid surrounding the developing fetus is a complex biological fluid rich in metabolically active bio-factors. The presence of extracellular vesicles (EVs) in amniotic fluid has been mainly related to fetal urine. We here characterized EVs from term amniotic fluid in terms of surface marker expression using different orthogonal techniques. EVs appeared to be a heterogeneous population expressing markers of renal, placental, epithelial and stem cells. Moreover, we compared amniotic fluid EVs from normal pregnancies with those of preeclampsia, a hypertensive disorder affecting up to 8% of pregnancies worldwide. An increase of CD105 (endoglin) expressing EVs was observed in preeclamptic amniotic fluid by bead-based cytofluorimetric analysis, and further confirmed using a chip-based analysis. HLA-G, a typical placental marker, was not co-expressed by the majority of CD105^+^ EVs, suggesting their origin from amniotic fluid cells. At a functional level, preeclampsia-derived EVs, but not normal pregnancy EVs, showed an antiangiogenic effect, possibly due to the decoy effect of endoglin. In addition, several miRNAs were differentially expressed in preeclampsia-derived EVs and directly related to the modulation of angiogenesis and trophoblast function. Our results provide a characterization of term amniotic fluid-EVs, supporting their origin from fetal and placental cells. In preeclampsia, the observed antiangiogenic characteristics of amniotic fluid-EVs may reflect the hypoxic and antiangiogenic microenvironment and could possibly impact on the developing fetus or on the surrounding fetal membranes.

## Introduction

Amniotic fluid is a complex biological fluid surrounding the developing fetus and in close proximity to the placenta (Underwood, Gilbert and Sherman, 2005). Its composition, after fetal skin keratinization, mainly consists of fetal cardiovascular, renal, pulmonary, and endothelial secretions (Roubelakis, Trohatou and Anagnou, 2012). Amniotic fluid has a relevant role in fetal physiology; it not only acts as a protective cushion to mechanical injury but also contains a plethora of active factors, including nutrients, growth and antimicrobial factors, cells and extracellular vesicles (EVs) (Underwood, Gilbert and Sherman, 2005).

EVs are a highly heterogeneous population of small particles enclosed in a lipid bilayer, released by almost all cell types, and involved in cell-to-cell signalling. In pregnancy, EVs are essential for diverse physiological and pathological processes (Jankovičová *et al*., 2020) and may play a role in angiogenesis and successful fetal development (Gebara *et al*., 2021). The role of EVs derived from placental trophoblasts has been studied extensively, as they are continuously released into the maternal bloodstream and play a crucial role in regulating the maternal immune response and pregnancy adaptation (Nair and Salomon, 2018). At variance, the profile and function of EVs present in amniotic fluid have been poorly investigated, as their characteristics have been mainly attributed to fetal urine production (Keller *et al*., 2007). Indeed, previous studies showed that a large proportion of EVs within the amniotic fluid express typical renal markers, such as aquaporin-2 and CD24, demonstrating their fetal urine origin (Keller *et al*., 2007). However, multiple cells may release EVs into the amniotic fluid, including the amnion, placental tissues and fetal cells (Dixon *et al*., 2018; McMaster *et al*., 1998). Interestingly, isolated amniotic epithelial or stem cells in culture were shown to release EVs with trophic, immunomodulating and anti-inflammatory functions (Balbi *et al*., 2017; Sedrakyan *et al*., 2017). It is therefore conceivable that EVs released from those cells are also present in the amniotic fluid. Moreover, a recent trial reported the safe administration of a pharmaceutical product, including term amniotic fluid-EVs, against the COVID-19 related cytokine storm (Mitrani *et al*., 2021). The analysis and characterization of amniotic fluid EVs is therefore of interest.

Preeclampsia is a complex pregnancy disorder characterized by new-onset hypertension with associated maternal organ dysfunction, including impaired liver function, renal damage (significant proteinuria, oligoanuria, increased creatinine), low platelets, haemolysis, decreased sPO_2_ or pulmonary edema, or visual and cerebral disturbances (Brown *et al*., 2018). Preeclampsia affects up to 8% of pregnancies world-wide (Duley, 2009) and causes significant maternal and perinatal morbidity and mortality, as well as long-term complications (Amaral *et al*., 2015). Abnormal placentation and insufficient trophoblast invasion are considered the main causes of preeclampsia (Kaufmann, Black and Huppertz, 2003), leading to the release of antiangiogenic factors that subsequently promote maternal endothelial dysfunction (Staff *et al*., 2007). Several studies highlighted the presence of altered EVs in the maternal circulation of preeclamptic women and their possible contribution to endothelial dysfunction (Escudero *et al*., 2016). Interestingly, EVs carrying antiangiogenic factors, such as soluble vascular endothelial growth factor receptor 1 (sFlt-1) and soluble endoglin, were found at high levels in the maternal circulation and were shown to be released by placental explants (Tannetta *et al*., 2013; Chang *et al*., 2018; Nair *et al*., 2021). Therefore, it is of interest to evaluate whether amniotic fluid EVs in preeclamptic pregnancies might have phenotypic or functional differences with respect to those of normal pregnancies.

In the present study, we aimed to characterize amniotic fluid-derived EVs from term normal and preeclamptic pregnancies. Surface marker expression was analyzed at a single-EV level using different techniques, such as super-resolution microscopy, ExoView array and bead-based cytofluorimetric analysis. In addition, their differential microRNA (miRNA) profile and angiogenic properties were investigated.

## Materials and methods

### Collection of amniotic fluid

Amniotic fluid samples (40 from normal pregnancies and 7 from preeclamptic pregnancies) were collected during caesarean delivery in the Department of Surgical Sciences at the University of Turin, after approval by the Ethics Review Board of the Health and Science City of Turin, (Città della Salute e della Scienza di Torino, protocol N° 0079734, CS2/320) and at the Maternity Department of St George’s University Hospital in London, after approval by the Local Ethics Committee (19/LO/0794) (Table 1). All patients provided preoperative written informed consent. The fluid was collected by sterile acupuncture after myometrial incision, before the incision of the amniotic membranes. Three of the preeclamptic patients (n=7) were affected by comorbidities, including fetal growth restriction, type two diabetes, and systemic lupus erythematosus. Further clinical information about preeclamptic patients can be found in Table 2.

**Table 1.** Clinical information showing the mean ± SD of gestational age, maternal age at delivery and the volume of the amniotic fluid obtained from normal and preeclamptic pregnancies.

|  | Normotensive pregnancies<br>(n=40) | Preeclampsia<br>(n=7) |
| --- | --- | --- |
| Gestational age (weeks) | 37.9±1.8 | 36.1±2.6 |
| Maternal age (years) | 35.7±5.2 | 32.8±6.7 |
| Amniotic fluid volume (ml) | 19.9±7.8 | 18.4±11.3 |

**Table 2.** Clinical information on individual preeclamptic patients showing the gestational age, maternal age at delivery, week of preeclampsia diagnosis, sFlt-1/PIGF ratio, other pathologies, treatment, and additional relevant information. PIGF:Placental growth factor.

|  | Age of gestation (weeks+days) | Maternal age (years) | Preeclampsia diagnosis (weeks) | sFlt-1/PIGF ratio | Other pathologies | Treatments | Other |
| --- | --- | --- | --- | --- | --- | --- | --- |
| 1 | 36+1 | 42 | 36 | 280 | - | Betamethasone<br>Nifedipine | Assisted reproduction techniques |
| 2 | 32+1 | 32 | 31 | 531 | - | Betamethasone<br>Nifedipine<br>MgSO<sub>4</sub> | Fetal growth restriction |
| 3 | 39+2 | 22 | 36 | 42 | - | - | - |
| 4 | 36+6 | 39 | 35 | - | Chronic hypertension, systemic lupus erythematosus | Methyldopa, Nifedipine, Hydroxychloroquine, Aspirin, Low Molecular Weight Heparin | - |
| 5 | 38+2 | 28 | 36 | - | Type 2 diabetes | Metformin, Insulin | - |
| 6 | 36+1 | 35 | 36 | - | - | Labetalol, Nifedipine | Fetal growth restriction |
| 7 | 36+2 | 33 | 33 | - | - | Labetalol, Nifedipine, Metformin, Insulin | - |

### Isolation and culture of term amniotic fluid mesenchymal stromal cells (AFSCs)

Term AFSCs were previously obtained and characterized, as described (Iampietro *et al*., 2020). Briefly, amniotic fluid was transferred into a sterile 50 mL falcon and centrifuged at 500 g for 10 minutes at room temperature. The cell pellet was resuspended in α-MEM Medium (Gibco/BRR, ThermoFisher, MA, USA) supplemented with 20% Chang Medium B (Irvine Scientific, Santa Ana, California, USA) and 2% Chang Medium C (Irvine Scientific), 20% Fetal Calf Serum (Invitrogen, Carlsbad, CA, USA), 50 IU/mL penicillin, 50 g/mL streptomycin, 5mM glutamine (all from Sigma-Aldrich, St. Louis, MO, USA). The cells were placed into a T25 flask, incubated at 37°C on 5% CO_2_ until the cells formed clusters (1-2 weeks). Afterwards, the medium was replaced twice a week until the cells reached confluency. Under these culture conditions, AFSCs expressed typical mesenchymal surface markers and differentiation properties, as previously described (Iampietro *et al*., 2020).

### EV Isolation from amniotic fluid

Amniotic fluid samples were diluted in PBS up to a volume of 20 ml and subjected to differential centrifugations. For the isolation of cells, samples underwent a 5 min centrifugation at 300 x g, at room temperature. A second centrifugation of 10 min was performed at 500 x g, at room temperature, for removal of debris. The supernatant was then passed through a cell sieve and underwent a 30 min ultracentrifugation (70 Ti rotor, Beckman Coulter, CA, USA) at 10,000 x g, at 4°C. The resulting pellet was discarded, and the supernatant was further centrifuged for 2 h at 100,000 x g at 4°C. The final step included resuspension of EV pellets in 1 ml 1% DMSO/PBS and additional filtration through a 0.22 µm filter. The recovered EVs were stored at −80°C.

### EV Isolation from AFSCs

The term AFSCs were grown to 70-80% confluency, washed twice with PBS and placed in serum free RPMI medium overnight. After 16 h of incubation, the RPMI medium was recovered and centrifuged at 300 x g for 5 min, 10,000 x g for 30 min, followed by ultracentrifugation at 100,000 x g for 2 h at 4°C. The EV pellet was resuspended in 1% DMSO/PBS and stored at −80°C.

### Smart SEC purification

Size-exclusion chromatography was applied to samples destined for miRNA arrays and ExoView analysis. Following differential ultracentrifugation, EV pellets were resuspended in 100 μl of PBS and applied to Smart SEC columns (SSEC100A-1, BioNova, Madrid, Spain). The samples were then processed as recommended by the manufacturer’s protocol.

### Transmission electron microscopy

One drop of PBS (about 25 µl) containing EV at 5.8×10^8^ was placed on a 400-mesh holey film grid; after staining with 2% uranyl acetate (for 2 minutes) the sample was observed with a Tecnai G2 transmission electron microscope (FEI Company, Hillsboro, OR, USA) operating at 100 kV. Images were captured with a Veleta digital camera (Olympus Soft Imaging System, Tokyo, Japan).

### NanoSight analysis

The size and concentration of EVs was determined by nanoparticle tracing analysis (NTA) performed by NanoSight LM10 system (Malvern Panalytical, Malvern, UK). Each vesicle sample was diluted (1:100) in PBS (filtered with a 0.22 µm filter). For each sample, a syringe pump flow of 30 was applied. Three videos of 60 seconds each were recorded and analyzed, calculating an average number of EV size and concentration (particles/ml). All samples were characterized with NTA 3.2 analytical software. The NTA settings were kept constant between the samples.

### Super-resolution microscopy

Super-resolution microscopy pictures of EVs were obtained using a temperature-controlled Nanoimager S Mark II microscope from ONI (Oxford Nanoimaging, Oxford, UK) equipped with a 100x, 1.4NA oil immersion objective, an XYZ closed-loop piezo 736 stage, and 405 nm/150 mW, 473 nm/1 W, 560 nm/1 W, 640 nm/1 W lasers and dual or triple emission channels split at 640 / and 555 nm. For sample preparation, 10 μl of 0.01% Poly-L-Lysine (Sigma-Aldrich, St. Louis, MO, USA) was placed on high-precision coverslips previously cleaned in a sonication bath twice in dH_2_O and once in KOH, in silicon gasket (Grace Bio-Labs, Sigma-Aldrich). The coverslips were placed at 37°C in a humidifying chamber for two hours. Excess of Poly-L-Lysine was removed. 1 μl of EVs (1×10^10^) resuspended in 9 μl of blocking solution (PBS-5% Bovine Serum Albumin) was pipetted into a previously coated well to attach overnight at +4°C. The next day, the sample was removed, and 10 μl of blocking solution was added into the wells for 30 min. Then, 2.5 μg of purified mouse anti-CD9 conjugated with Atto 488 dye (ONI), anti-CD63 antibodies (SC-5275, SC-31234, Santa Cruz, CA, USA) conjugated with Alexa Fluor 647 dye, HLA-G antibody (SC-21799, Santa Cruz) conjugated with Alexa Fluor 555 dye, were prepared using the Apex Antibody Labelling Kit (Invitrogen) according to the manufacturer’s protocol. One μl of each antibody was added into the blocking buffer at a final dilution of 1:10. Samples were incubated in the dark at +4°C. The day after, samples were washed twice with PBS, and 10 μl of the mixed ONI B-Cubed Imaging Buffer (Alfatest, Rome, Italy) was added for amplification of the EV signalling. Before each imaging session, bead slide calibration was performed for aligning the channels, to achieve a channel mapping precision smaller than 12 nm. Images were taken in dSTORM mode using 30% laser power for the 647 nm, 50% laser power for the 488 nm laser, and 30% for the 555 channel. Two-channels (647 and 555) dSTORM data (5000 frames per channel) or three-channels (2000 frames per channel) (647, 555 and 488) were acquired sequentially at 30 Hz (Hertz) in total reflection fluorescence (TIRF) mode. Single-molecule data was filtered using NimOS software (v.1.18.3, ONI) based on the point spread function shape, photon count and localization precision to minimize background noise and remove low-precision and non-specific colocalization. Data has been processed with the Collaborative Discovery (CODI) online analysis platform www.alto.codi.bio from ONI and the drift correction pipeline version 0.2.3 was used. Clustering analysis was performed on localizations and BD clustering constrained parameters were defined (photon count 300-max, sigma 0-200 nm, p-value 0-1, localization precision 0-20 nm). Colocalization was defined by a minimum number of localizations for each fluorophore/protein within a distance of 100 nm or distance used from the centroid position of a cluster.

### Western Blot

For protein analysis, EVs were lysed at 4°C for 30 min in RIPA buffer (20 nM Tris-HCl, 150 nM NaCl, 1% deoxycholate, 0.1% SDS, 1% Triton X-100, pH 7.8) supplemented with a cocktail of protease and phosphatase inhibitors and PMSF (Sigma-Aldrich). For protein concentrations of less than 2 mg/ml, a precipitation step was applied. The pellet was resuspended in Laemmli Buffer and the sample was loaded onto the gel. As determined by the Bradford method, aliquots of the cell lysates containing 10-15 µg of proteins were run on 4-20% acrylamide gel SDS-PAGE under reducing conditions. The transfer of proteins onto a PVDF membrane was performed using the iBlot™ Dry Blotting System (Life Technology). Primary antibodies used were: CD81 at 1:200 (SC-31234), CD63 at 1:200 (SC-5275), HLA-G at 1:200 (SC-21799), (all from Santa Cruz), CD105 at 1:1000 (362-820, Ancell) and at 1:2000 calreticulin (2891S, Cell Signaling Technology, Danvers, MA, USA). Chemiluminescent signal was detected using the ECL substrate (Bio-Rad), and bands were detected by the ChemiDoc system (Bio-rad, Hercules, CA, USA).

### MACSPlex

EV samples were tested with a bead-based multiplex analysis (MACSPlex exosome kit, human, Miltenyi Biotec, Bergisch Galdbach, Germany). EV samples were diluted with MACSPlex buffer to a final volume of 120 µl with 5.8×10^8^ EVs per sample. Fifteen µl of MACSPlex Exosome Capture Beads and 15 µl of the antibody cocktail composed of APC conjugated CD9, CD63 and CD81 were added to the buffer containing EVs and incubated on an orbital shaker at 12 rpm for 1 hour protected from light, at room temperature. After the incubation, 1 ml of MACSPlex buffer was added and the samples were centrifuged at 3,000 g for 5 min. After a second wash, the samples were incubated for a further 15 min on the orbital shaker. Thereafter the samples were centrifuged and prepared for flow cytometry analysis. Median fluorescence intensity was calculated for all 39 bead populations. The analysis of the samples was performed by subtraction of the control and the fluorescence controls. For analysis of AFSC-EVs, the same method was followed with the substitution of the fluorescent antibody mix against tetraspanins (CD63, CD81 and CD9) with the APC labelled HLA-G antibody (130111852, Miltenyi Biotec, same volume of 15 µl). This method ensured that EVs attached to the Ab conjugated beads were only detected if positive for HLA-G.

### ExoView array

For the analysis with the ExoView platform™ (NanoView Biosciences, Boston, MA, USA), EVs were further purified with Smart SEC columns (BioNova), and the collected fractions were applied directly to the ExoView chips. 10-35 µl of the sample with a concentration of 5×10^8^ EVs was applied to a final volume of 35 µl in the incubation solution. EVs were then incubated overnight at room temperature. Immuno-fluorescence staining was performed using antibodies provided by the company (0.6 µl of CD63, CD81, CD9). Alexa Fluor 488 conjugated anti-CD117 (ab2164459, Abcam, Cambridge, UK), anti-Tie-2 (ab24859, Abcam), anti-CD105 (362-820, Ancell) and HLA-G conjugated with Alexa Fluor 555 (SC-21799, Santa Cruz) were used for detection of EV subpopulations. For all antibodies, the Apex Antibody Labelling Kit was used (Invitrogen). All antibody conjugations were performed according to manufacturer’s protocol. Chips were analyzed using the ExoView™ R100 reader using ExoView Scanner software (v 3.0). The data were analyzed using ExoView Analyzer (v 3.0). Two sets of ExoView chips were tested. The first set of chips were pre-spotted with CD63/CD81/CD9, with addition of fluorescent antibodies CD63, CD81, CD9. The second set of chips were pre-spotted with CD63/CD81/CD9 with the addition of fluorescent antibodies HLA-G, CD117, Tie-2 and CD105.

### Angiogenesis

Human umbilical vascular endothelial cells (HUVEC) were bought from ATCC (ATCC-PCS-100-010, Manassas, VA, USA). Cells were cultured until the 6th passage in complete EBM medium (Lonza, Basel, Switzerland). Cell medium was changed three times per week, and cells were passaged when they reached 70-80% confluency. The cells were split by washing in PBS, then applying 2 ml of 10% trypsin-EDTA and re-plated on T75 flasks coated with attachment factor (S-006-100, Gibco). *In vitro* formation of capillary-like structures was performed on growth factor–reduced Matrigel (356231, Corning, NY, USA). HUVEC cells were mixed with EVs (1000 EVs/cell) and/or anti-CD105 Ab TRC-105 8 µg/ml (TRACON Pharmaceuticals, CA, USA) and recombinant soluble CD105 (s-Endoglin, 100 ng/ml; C-60059, PromoKine, Heidelberg, Germany) in complete EBM medium and seeded at a density of 15×10^3^ cells per well on a 24-well plate. For combination of EVs with the anti-CD105 Ab TRC-105, both components were mixed and incubated for 25 min at room temperature, prior to adding them to the cell suspension. Cells were periodically observed with a Nikon TE2000E inverted microscope (Nikon, Tokyo, Japan), and experimental results were recorded after 16 h; 5 images were taken per well. Image analysis was performed with the ImageJ software v.1.53c, using the Angiogenesis Analyzer. The data from three independent experiments were expressed as the mean ± SD of tube length in arbitrary units per field.

### miRNA array analysis

Total EV RNA was extracted from 3 normal and 3 preeclamptic amniotic fluid samples after ultracentrifugation and SEC. All samples were processed separately. RNA was extracted using *mir*Vana miRNA Isolation Kit (AM1560, ThermoFisher) following the manufacturer’s protocol. The RNA concentration was measured using NanoDrop (NanoDrop ND-1000, ThermoFisher) and precipitated to a final concentration of 10 ng/μl. The total cDNA was pre-amplified prior to miRNA quantification. Expression of 754 different miRNAs was evaluated by quantitative real-time PCR using TaqMan Array MicroRNA card A (v.3.0) and B (v.3.0) and according to Megaplex manufacturers protocol. All the samples were run separately. Quantification was performed using a QuantStudio 12K Flex Real-Time PCR System (Applied Biosystems, Waltham, MA, USA). Ct values were analyzed by Expression Suite software v1.3 using global normalization. A maximum threshold of 35 was applied for analysis of EVs from normal pregnancies in comparison to preeclampsia.

### Quantitative real-time PCR

Validation of the miRNAs was performed by quantitative real-time PCR (qRT-PCR) on 3 different amniotic fluid EV samples for normal pregnancy and preeclampsia. The miRNAs were reverse-transcribed into cDNA using a High Capacity cDNA reverse transcription kit (Applied Biosystems) following the manufacturer’s protocol. The qRT-PCR was performed using 20 miRNA primers (Supplementary Table 1) and miScript SYBR Green PCR Kit (Qiagen, Hilden, Germany). RNA U6 was used as the endogenous control. The samples were analysed using the comparative CT method. In total 20 miRNAs were selected and validated, based on literature research via Pubmed interrogation with the following keywords: “miRNAx” and “angiogenesis or placenta or pre-eclampsia” or “miRNAx” and “angiogenesis or placenta or pre-eclampsia” and extracellular vesicles”.

### Acquisition of PE-related miRNAs gene targets and transcription factors and network visualization

We obtained PE associated miRNAs based on previously mentioned experiments. miRTarBase v8.0 was used to identify gene targets and extract miRNA-gene pairs (Huang *et al*., 2020). To minimize false positives, only strong-evidence miRNA-target pairs were considered. Literature-curated transcription factor (TF)-miRNA pairs of deregulated miRNAs were extracted from TransmiR (Tong *et al*., 2019) miRNA, target gene, and TF interaction pairs were visualized in Cytoscape v3.8.2 (Shannon *et al*., 2003). The interactions between genes and TF, and interacting genes were integrated using the Cytoscape App Bisogenet v3.0.0 (Martin *et al*., 2010).

### Network enrichment analysis

The analysis of overrepresented gene ontology (GO) biological process terms and pathways associated with the target genes, was performed using EnrichR webtools (Chen *et al*., 2013). Ontology and pathway GO terms with an adjusted p-value <0.05 were considered significantly overrepresented. In addition, the Cytoscape App BinGo was used to visualize and cluster the gene annotation terms into groups (Maere, Heymans and Kuiper, 2005). BinGo results were corrected for multiple testing using Benjamini-Hochberg false discovery rate correction. The KEGG, Reactome and WikiPathways databases, as well as the Tissue Expression ArchS4 databases, were used to obtain specific gene annotations (Ogata *et al*., 1999; Rouillard *et al*., 2016; Jassal *et al*., 2020). The comparison of miRNA-gene target interactions and GO biological processes to previous studies (Pillay *et al*., 2019 and Salomon *et al*., 2017) was performed using Cytoscape ClueGO plugin (Bindea *et al*., 2009). GO biological process function terms with an adjusted p value <0.001 after Bonferroni step down correction and term fusion were considered.

### Synergistically working miRNAs

MiRNAs with binding sites near each other (between 13-35 nucleotides) can cooperate in target repression. We identified cooperating miRNA pairs of upregulated preeclampsia-associated miRNAs using the TriplexRNA database (https://triplexrna.org/). The results were filtered for targets involved in angiogenesis based on GO terms (Lai *et al*., 2012).

### Statistical tests

All statistical analyses were performed using GraphPad Prism software (v. 8.00; GraphPad, CA, USA). For comparisons of each group, an unpaired Student’s t-test was used. FWelch’s correction was performed when the variance across groups was assumed to be unequal. Differences were considered significant when the p-value was <0.05. For multiple comparison analyses, ANOVA with Bonferroni multicomparison post hoc test. A two-tailed *p*-value <0.05 was considered statistically significant.

## Results

### Characterization of amniotic fluid-derived EVs

EVs were isolated from term amniotic fluid of 40 normal pregnancies (Table 1) by differential ultracentrifugation followed by filtration and, for selected experiments (miRNA and ExoView analysis), by size-exclusion chromatography. The normal pregnancy EVs (NP-EVs) were first characterized based on their size and concentration using nanoparticle tracking analysis (Figure 1A). Homogeneous concentration and particle size were obtained among the different samples, with a mean size of 221.8±6.2 nm. Western Blot analysis confirmed the expression of EV specific markers (CD63, CD81) and revealed the presence of the placental maker HLA-G, previously shown to characterize placental-derived EVs (Orozco *et al*., 2009) (Figure 1B). The absence of the cytoplasmic marker calreticulin showed a lack of cellular contamination (Figure 1B). TEM showed heterogeneous EV populations (particles ranging from approximately 50 to approximately 200 nm in diameter) (Figure 1C). To quantify the expression of CD63 and HLA-G and to assess their possible colocalization, we performed super-resolution imaging of NP-EVs using dSTORM. More than 6,000 single-EV images were acquired and analyzed. Clustering analysis showed that the majority (around 87%) of CD63^+^ EVs also expressed HLA-G (Figure 1D).

**Figure 1.**
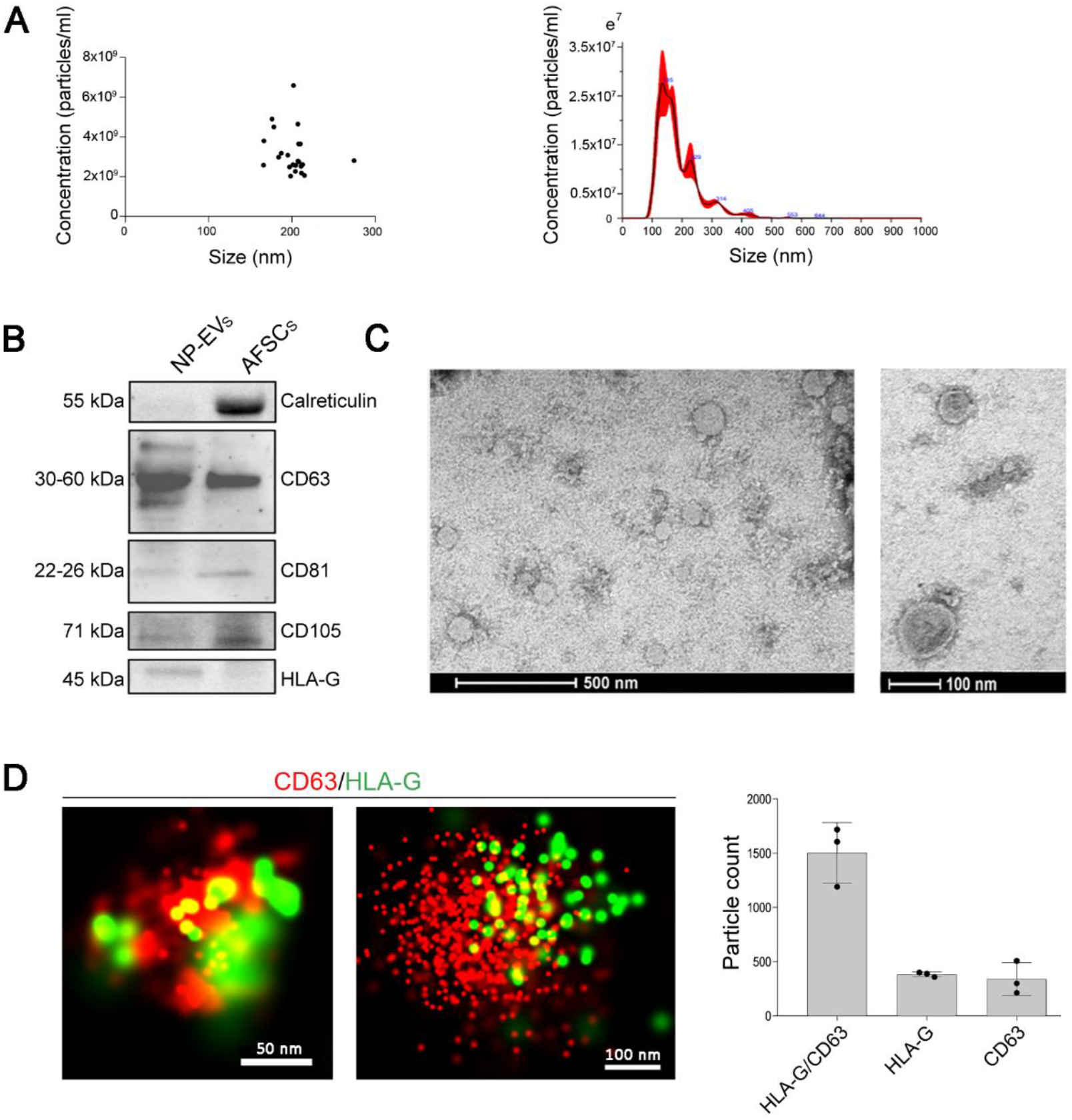
Characterization of NP-EVs. **A)** Nanoparticle tracking analysis of NP-EVs (n=30) showing homogenous size and concentration. Dilution of 1:100 in PBS in 1ml was used to test the samples (left panel). A representative graph showing size distribution of a NP-EV sample (right panel). **B)** Representative Western Blot image showing the presence of CD63, CD81, CD105 and HLA-G in NP-EVs (10-15 µg proteins) and in AFSC lysate (10-15 µg proteins) and the presence of calreticulin in cell lysate only. Three different NP-EV samples were tested with similar results. **C)** Representative transmission electron microscopy images at low- and high-power fields of NP-EVs showing heterogenous EV population. **D)** Representative super-resolution microscopy images of single amniotic fluid-derived EVs expressing CD63 (red) and HLA-G (green). The number of single and double positive EVs for CD63 and HLA-G was analysed in 3 NP-EV preparations using the CODI software; the graph shows the mean ± SD of a cumulative analysis of 10 fields for each preparation, total EV number: 6,676.

In addition, super-resolution microscopy was used to analyze the size and expression of tetraspanin markers (CD63, CD81 and CD9) at a single-molecule level in NP-EVs (Figure 2). The results showed that NP-EVs were a heterogeneous population with highly variable tetraspanin expression (Figure 2E). The analysis of more than 20,000 EVs using a dedicated software (https://alto.codi.bio/) showed that the majority of EVs were either triple positive for CD63, CD81 and CD9 or double positive for CD81 and CD9, with lower fractions expressing double or single tetraspanins (Figure 2E). The overall average particle size evaluated by the super-resolution microscope was 121±26 nm. This measurement, based on specific EV detection using tetraspanins, overlaps the results obtained by transmission electron microscopy. The larger EV size detected by nanoparticle tracking analysis can be possibly explained by the evaluation of the EV hydrodynamic size, under solution using this technique, in comparison with the measurement of the dry radius with microscopy techniques (Montis *et al*., 2017). In addition, possible artifacts including EV aggregates or non-EV particles can be detected by nanoparticle tracking analysis, suggesting that it should be applied for size evaluation together with other side techniques, as previously described (Skovronova *et al*., 2021).

**Figure 2.**
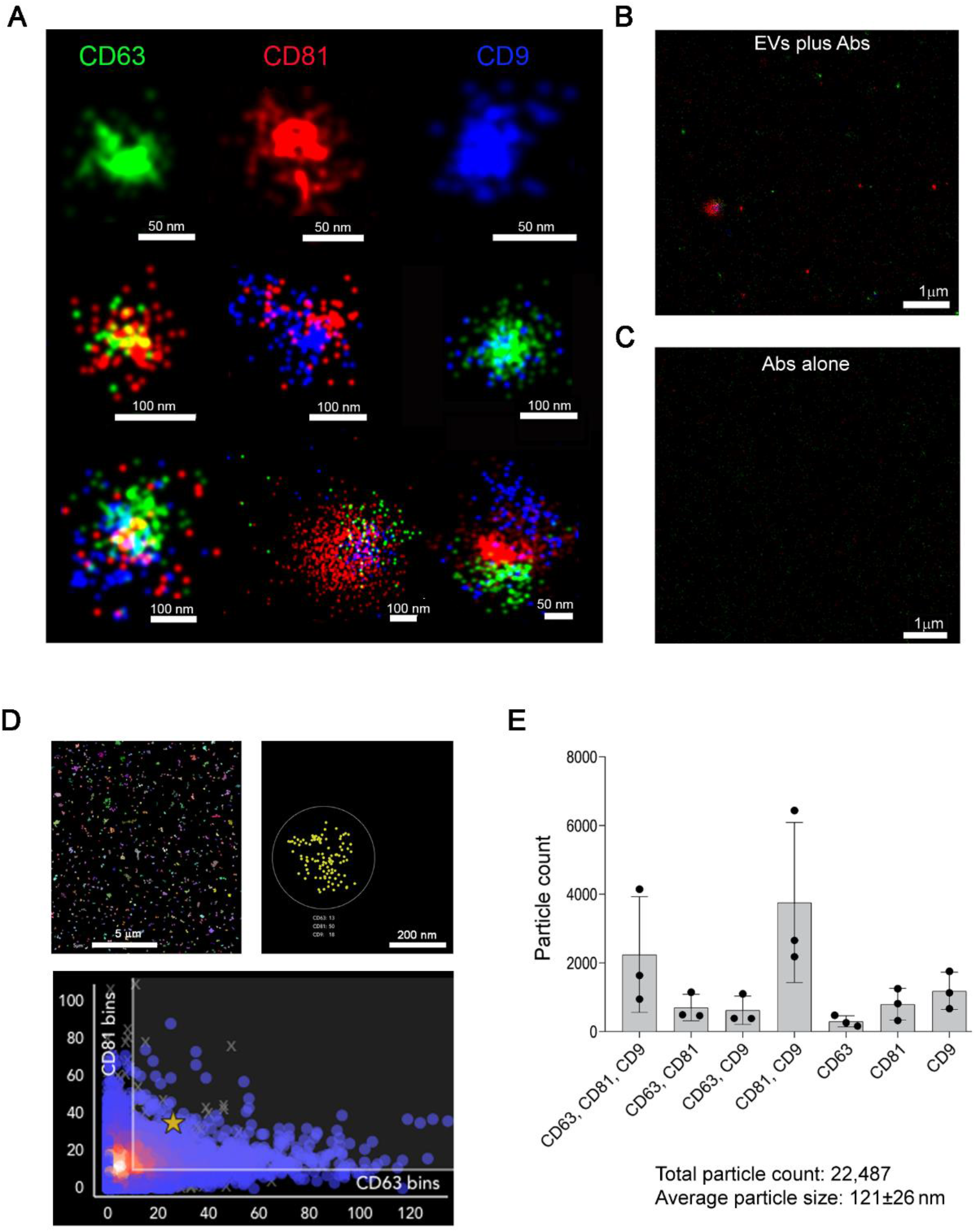
Super-resolution microscopy analysis of tetraspanins expression in NP-EVs. **A)** Representative super-resolution microscopy images of NP-EVs showing single, double, and triple expression of CD63 (green), CD81 (red), CD9 (blue). The corresponding scale bare is below each EV image. **B)** Representative super-resolution images, at low magnification, showing staining with anti-tetraspanin antibodies in the presence of EVs (EVs plus Abs) or **C)** in the absence of EVs (Abs alone). **D)** Representative clustering strategy of NP-EV analysis showing a large field of view (left panel), a selected cluster (right panel) and a graph (bottom panel) of CD81/CD63 cluster distribution. **E)** Clustering analysis of super-resolution microscopy images showing the single, double, and triple positive EV fractions expressing the tetraspanin markers. The analyses were performed in 3 NP-EV preparations using a CODI software; the graph shows the mean ± SD of a cumulative analysis of 10 fields for each preparation.

### Comparison of amniotic fluid EVs from normal and preeclamptic pregnancies

EVs were also isolated from term-amniotic fluid of 7 preeclamptic pregnancies (Table 1). The different volumes and EV concentrations in each sample are reported in Supplementary Table 1. No difference in mean EV concentration between NP-EVs and PE-EVs (2.85×10^9^ and 2.17×10^9^ particles/ml, respectively) was observed, although the relevance of this parameter is limited by the dynamic nature of this biofluid (Underwood, Gilbert and Sherman, 2005). As amniotic fluid contains EVs originating from several sources, a comprehensive analysis of EV surface receptors characterizing several cell types was performed using MACSPlex, a bead-based cytofluorimetric kit able to perform a semiquantitative fluorescent analysis of 39 different EV surface markers. Figure 3A shows the most relevant markers expressed. EVs within the amniotic fluid of both normal and preeclamptic pregnancies highly expressed tetraspanins, as confirmed with the methods shown above. In addition, the typical amniotic fluid marker, CD24 (Keller *et al*., 2007) and CD133, known to be highly expressed by urine EVs (Dimuccio *et al*., 2014) were present at a high level in both EV types, indicating the presence of EVs derived from fetal urine. Expression of the embryonic stem and epithelial cell marker CD326 was detected in both samples. Interestingly, a significantly higher expression of the progenitor/mesenchymal marker SSEA-4 and CD105 was detected in PE-EVs (Figure 3A). No expression of leukocyte (CD3, CD4, CD8, CD45, CD56, CD19), endothelial (CD31) or platelet (CD42a and CD49e) markers was observed (data not shown).

**Figure 3.**
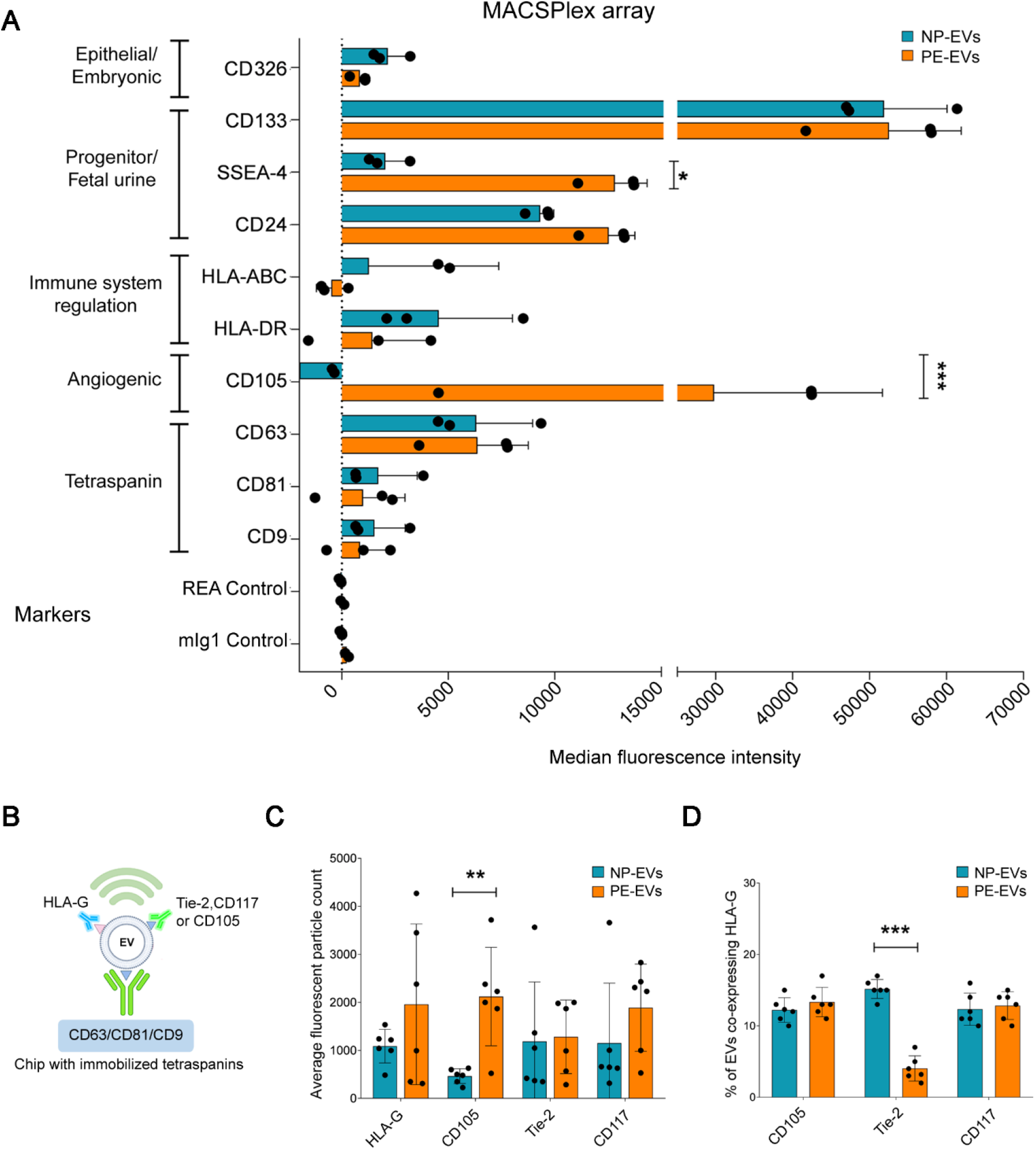
Increased CD105 expression in PE-EVs. **A)** MACSPlex analysis showing the median fluorescence intensity of surface markers characteristic of EVs (tetraspanins) or different cell of origin, expressed by NP- and PE-EVs (n=3). Expression of CD105 and SSEA-4 was significantly increased in PE-EVs. 5.8×10^8^ EVs were analysed, diluted to a final volume of 120 µl of MACSPlex buffer. **B)** Diagram explaining the experimental method behind ExoView technology in relation to the graphs in panel C. **C and D)** ExoView analysis of amniotic fluid-derived NP-EVs (n=6) and PE-EVs (n=6). **C)** Comparison of the expression of HLA-G, Tie-2, CD105 and CD117 (c-kit) shown as average fluorescent particle count in NP-EVs vs PE-EVs from combined tetraspanins capture of CD63, CD81 and CD9. **D)** Normalized expression of HLA-G positive EVs co-expressing other angiogenic (CD105 and Tie-2) and stem cell (CD117) markers. 5.8×10^8^ EVs in final volume of 35 µl of buffer were used for all samples. Unpaired Student’s t-test:*=P<0.05, * *=P<0.01, ***=P<0.001

ExoView analysis was subsequently used to assess at single-EV level the expression of NP-EV and PE-EV surface markers of interest. After EV affinity binding to tetraspanins, CD105, Tie-2, CD117 (c-kit) and HLA-G expression were evaluated on captured EVs (Figure 3C). The analysis confirmed the presence of HLA-G, as shown by super-resolution microscopy, and revealed the presence of the angiogenic marker Tie-2 and of the stem cell marker CD117 on both NP- and PE-EVs. In addition, the analysis confirmed the increased expression of CD105 in PE-EVs with respect to NP-EVs (Figure 3C). By simultaneously detecting HLA-G along with the other makers, we were able to assess their co-expression. The analysis showed that the HLA-G expressing EVs displayed similar CD105 and c-kit markers levels, suggesting that the increased CD105 expressing EVs present in PE were not of placental origin (Figure 3D). At variance, Tie-2 levels were significantly lower in placental EVs of PE pregnancies. No differences in tetraspanin levels, used as control, were observed (Supplementary Figure 1B). The increased CD105 expression in PE-EVs with respect to NP-EVs was also confirmed by Western Blot analysis (Supplementary Figure 2).

Finally, we tested the characteristics of EVs derived from cultured term AFSCs, as a possible source of amniotic fluid-EVs (Figure 4). Detection of AFSC-derived EVs captured on tetraspanin coated chips revealed the presence of distinct subpopulations of HLA-G and CD105 expressing EVs, being CD105^+^ EVs the larger fraction. No co-expression of the two markers was observed (Figure 4A and B). Analysis of the HLA-G^+^ AFSC-EV fraction by MACSPlex analysis confirmed the lack of CD105 expression (Figure 4C and D). These data suggest that the CD105^+^ EV fraction observed in PE-EVs could possibly derive from cells of fetal or fetal membrane origin, including AFSCs.

**Figure 4.**
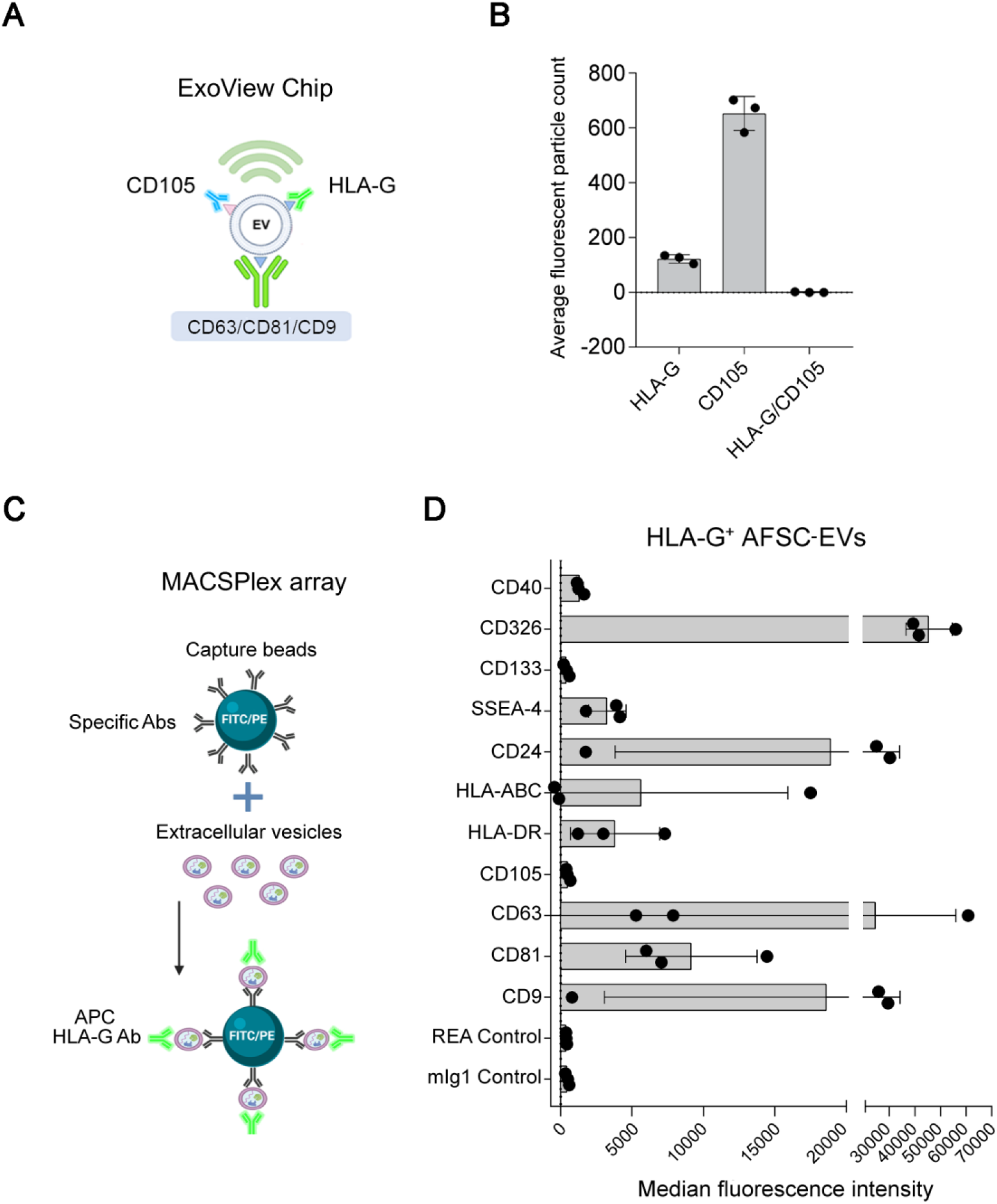
Characterization of HLA-G-expressing EVs from term AFSCs. **A)** Diagram explaining the experimental method in relation to the graph in panel B. **B)** Fluorescent particle count of AFSC-EVs captured on tetraspanin-coated chip analyzed by ExoView, showing expression of HLA-G^+^ and CD105^+^ EVs, but lack of co-expression of the markers. **C)** Graphical representation of MASCPlex analyses to characterize HLA-G^+^ AFSC-EVs, using Ab-coated fluorescent beads and APC-labelled anti-HLA-G Ab. **D**) The graph shows the median fluorescence intensity of surface markers co-expressed on HLA-G^+^ AFSC-EVs (n=3). AFSC-EVs (5.8×10^8^) were gated for the HLA-G positivity and analysed for the expression of a panel of markers. No expression of CD105 was detected. Illustrations in panels A and C were created with https://biorender.com.

### Functional characterization of NP-EVs and PE-EVs on angiogenesis

Soluble endoglin is a differentially spliced form of endoglin and is a relevant antiangiogenic factor in PE, acting as a decoy receptor for transforming growth factor beta (TGF-β) (Valluru *et al*., 2011). To test the angiogenic properties of NP-EVs, PE-EVs, and the possible role of CD105 increased expression on the EV surface, a tube formation assay was performed. Amniotic fluid-derived EVs both from NP and PE, showed a significant reduction in the organization of the HUVEC into capillary-like structures on Matrigel, with respect to the positive control, with the PE-EVs being significantly more effective than NP-EVs in reducing tube formation. To test the relative role of the increased expression level of CD105 on the surface of PE-EVs, TRC-105, an anti-CD105 monoclonal Ab used as an antiangiogenic drug (Brossa, Buono and Bussolati, 2018), was applied. We found that the addition of TRC-105 to PE-EVs, but not to NP-EVs, prior to HUVEC stimulation significantly abolished their inhibitory effect (Figure 5), indicating the role of surface CD105 in angiogenesis reduction.

**Figure 5.**
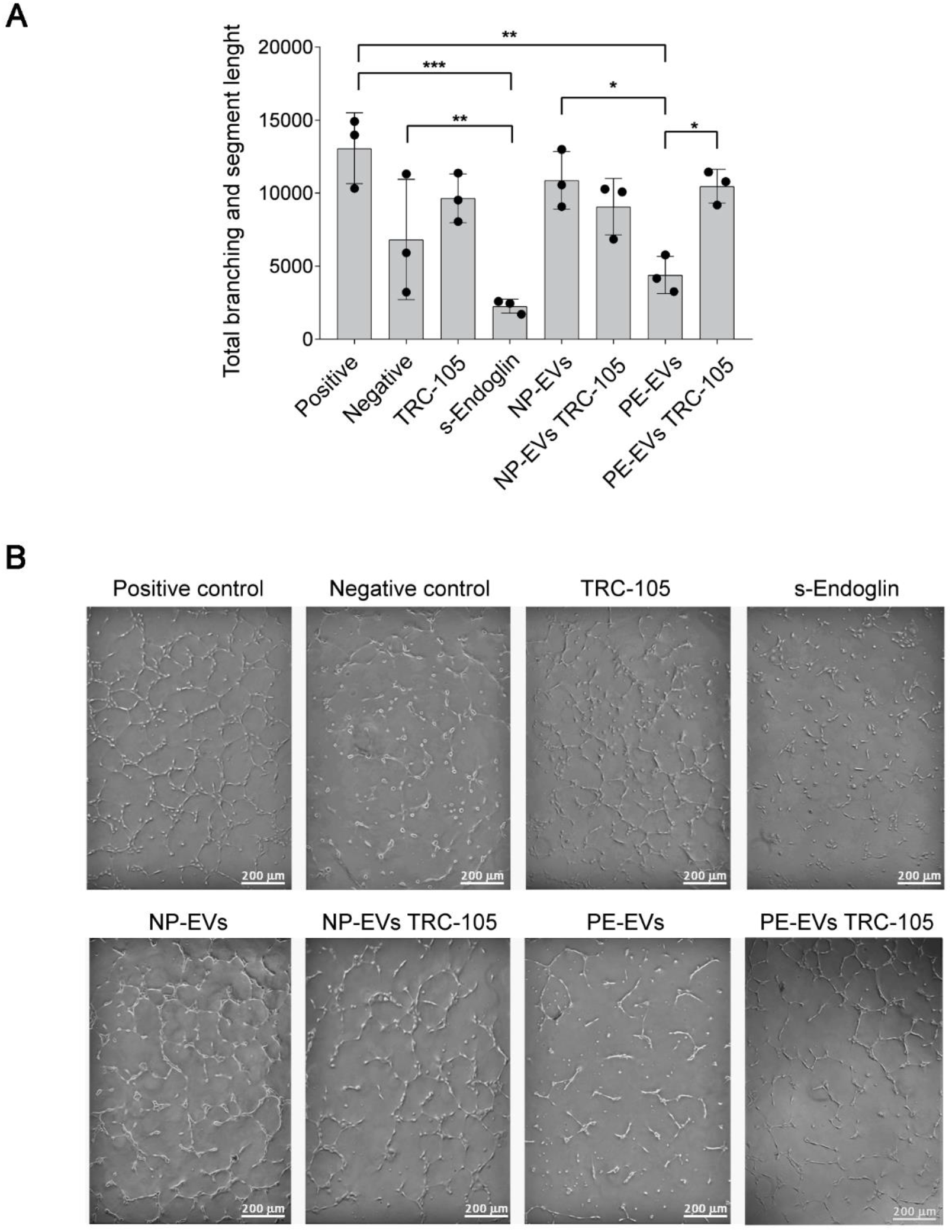
Effect of NP-EVs and PE-EVs on tube formation and role of CD105. **A)** Quantification of tube lengths of HUVEC treated with NP-EVs, PE-EVs with or without anti-CD105 Ab (TRC-105) (8 µg/ml). A concentration of 1000 EVs/cell was used. Positive control: complete EBM medium; negative control: EBM medium without FBS. s-Endoglin (100 ng/ml) was used as control for angiogenesis inhibition. Statistical analysis was performed using ANOVA with Bonferroni’s post-hoc test. *=P<0.05, **=P<0.01, ***=P<0.001. **B)** Representative images of the tube formation assay.

### Differential miRNA expression in NP-EVs and PE-EVs

In order to identify the key regulatory miRNA in amniotic-derived EVs and their possible differential expression in PE-EVs, we performed miRNA arrays on SEC-purified EV preparations from 3 NP and 3 PE amniotic fluids. A large number of miRNAs were present in amniotic fluid-EVs. The 3 sample repeats were combined for ease of interpretation. The raw data can be found at https://fairdomhub.org/projects/244, showing miRNA expression for each individual sample. The top and bottom 10 expressed miRNAs in NP-EVs are shown in Supplementary Table 2. Overall, 46 miRNAs were detected as differentially expressed in PE-EVs vs NP-EVs (Figure 6A and B). The complete list of miRNAs can be found at https://fairdomhub.org/data_files/4182?version=1. In particular, 20 miRNAs of interest were selected for qRT-PCR validation as previously described in EVs, biological fluids or placental tissue from preeclamptic women from literature analysis (Supplementary Table 4). Results confirmed a deregulated trend of the majority of miRNAs in PE-EVs, with let-7e, miR-15b, miR-195,miR-613, miR-26a and miR-144 showing significance (Figure 6C).

**Figure 6.**
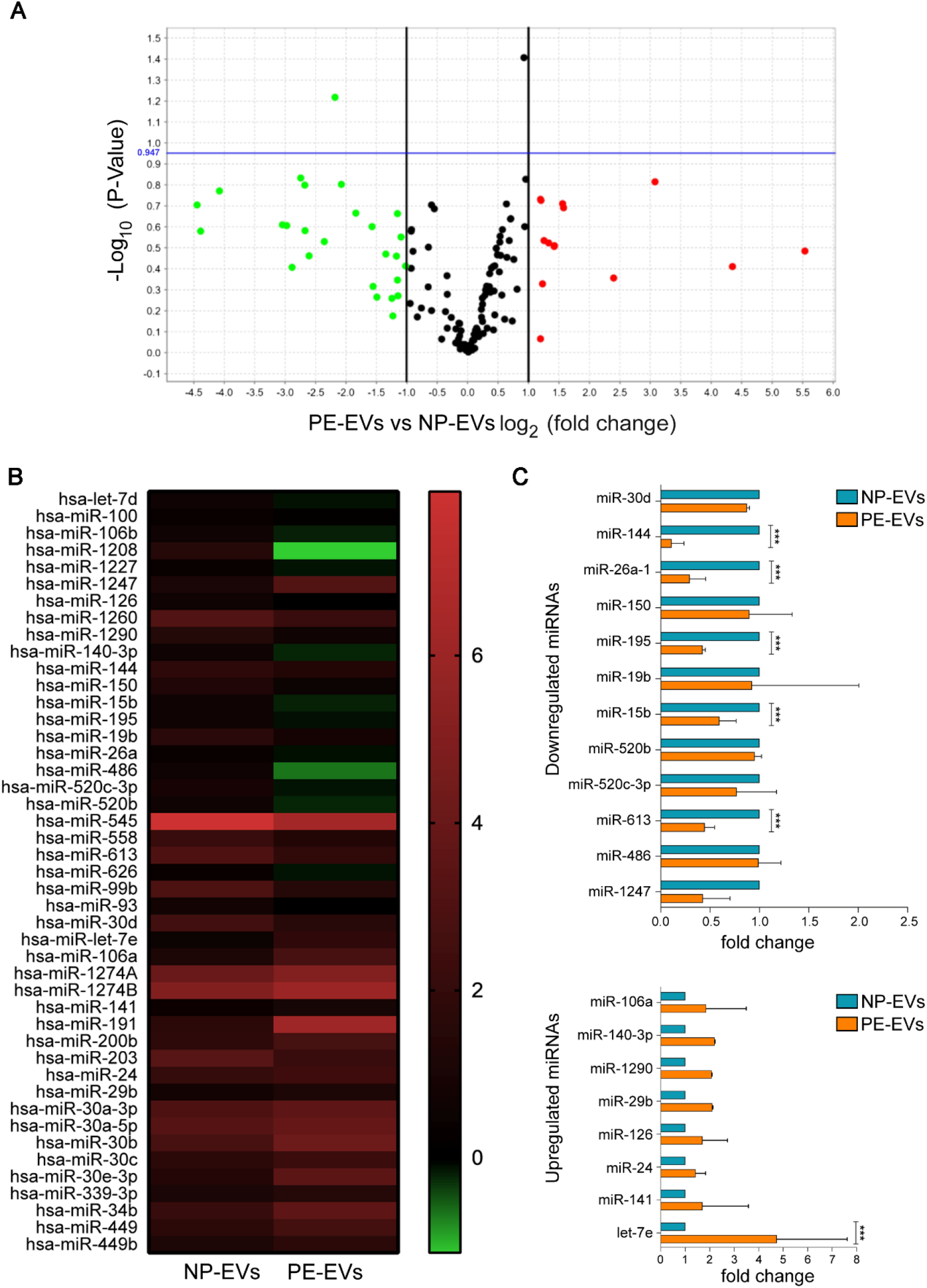
MiRNA analysis of NP-EVs and PE-EVs. **A)** Volcano plot showing the differential expression of miRNAs derived from NP-EVs vs PE-EVs. **B)** Heatmap showing the miRNAs with the highest fold changes between the samples. **C**) Validation of the deregulated miRNAs, normalized to RNU6 and CTL, as evaluated by qRT-PCR analysis. Data are expressed as mean ± SD of at least three different experiments normalized to CTL and to one, with let-7e, miR-195, miR-613, miR-15b, miR-26a-1, and miR-144, showing significant regulation differences. Unpaired Student’s t-test: ***=P<0.001.

### Bioinformatic analysis

The search of deregulated miRNA gene targets revealed more than 1000 strong evidence miRNA-gene interactions, involving 667 unique genes, with the vascular endothelial growth factor (VEGF) gene being central in the interactome (Supplementary Figure 3). Further functional enrichment analysis of GO terms and pathways supported the overall miRNA involvement in the regulation of angiogenesis and vascular remodeling (Supplementary Figure 4 and 6).

Accordingly, the analysis of target tissue and cell specific gene expression profile identified, among the top represented, myoblast, omentum and placenta (Supplementary Figure 5). Finally, analysing for miRNAs acting on a common target and resulting in an increased inhibitory effect *via* triplex analysis, COL22A1, coding for collagen XXII, appeared to be targeted by cooperative miRNAs with the lowest predicted repression (Supplementary Figure 6). Further details can be found in the bioinformatic section of supplementary data. The complete list of miRNA gene targets and network enrichment results from EnrichR and BinGo can be found at: https://fairdomhub.org/projects/244.

## Discussion

As reproductive EV research advances, investigation of placenta, surrounding gestational tissues, and amniotic fluid is of profound importance, in order to understand the factors involved in fetal development and maternal health. This paper provides for the first time a deep characterization of EVs present in term amniotic fluid in normal and preeclamptic pregnancies. Using several orthogonal techniques (chip-based platform, cytofluorimetric bead-based multiplex assay and super-resolution microscopy analysis), we show that EVs of term amniotic fluid represent a heterogeneous population of EVs of multiple origins, including placental tissues, fetal urine, and stem cells. Moreover, we identified potential antiangiogenic properties of amniotic fluid-EVs from preeclamptic pregnancies, supported by the specific upregulation of the CD105 surface marker and the deregulation of several angiogenic miRNAs (Figure 7).

**Figure 7.**
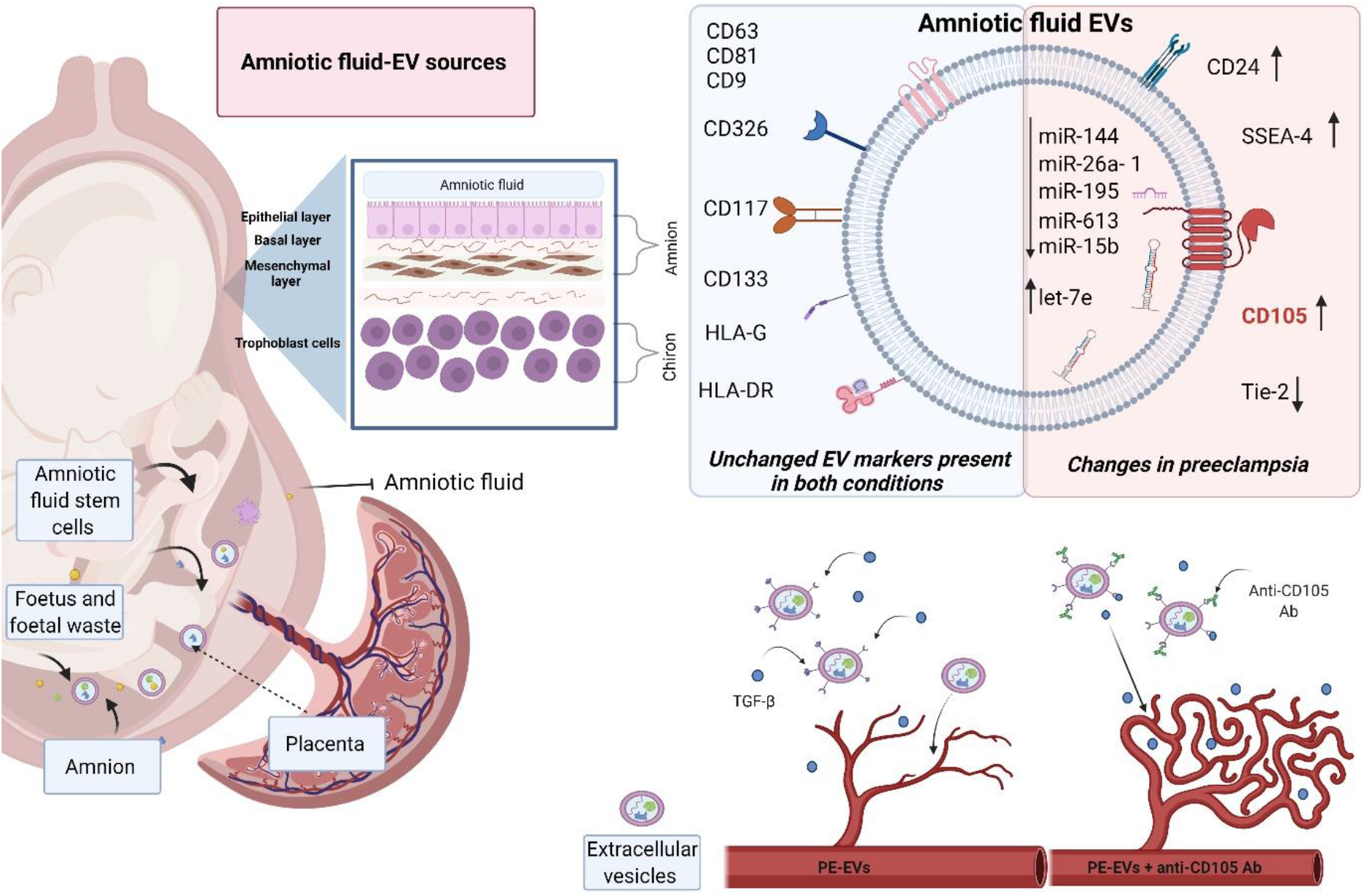
Graphical illustration summarizing the main findings. The figure shows the multiple possible sources of term amniotic fluid-derived EVs and the main changes in PE-EVs. Their antiangiogenic properties are supported by the specific upregulation of CD105 surface marker and the deregulation of several angiogenic miRNAs. The image was created with https://biorender.com.

We here characterized EVs isolated from term amniotic fluid using sub-sequential centrifugation, and we further purified them for the identification of specific surface receptors and miRNA cargo using size-exclusion chromatography to remove contaminating factors possibly present in the amniotic fluid (Monguió-Tortajada *et al*., 2019; Théry *et al*., 2018). The characterization of NP-EVs at a single-EV level revealed a highly heterogeneous population of EVs with variable tetraspanins expression. These results are in agreement with those reported by Han and colleagues on colocalization analysis of tetraspanins in EVs derived from different cell lines, showing distinct fractions of single, double or triple co-expressing EVs (Han *et al*., 2021).

Moreover, based on surface marker expression as well as miRNA content, we found that EVs within the term amniotic fluid may originate from various sources. Indeed, amniotic fluid-EVs expressing CD24 were mainly derived from fetal urine (Keller *et al*., 2007). In this study, we confirmed the possible cellular origin of the EVs attributed to fetal urine by expression not only of CD24 but also of CD133, which is also considered to be highly present in urinary EVs (Dimuccio *et al*., 2014). Additionally, amniotic fluid-EVs expressed mesenchymal and stem markers, possibly suggesting the origin from CD117^+^ amniotic fluid stem cells (Cananzi, Atala and De Coppi, 2009), amniotic CD105^+^ mesenchymal stem cells (Baghaei *et al*., 2017), and SSEA-4^+^ expressing embryonic stem cells (Noisa *et al*., 2012). Interestingly, we also showed that a large fraction of amniotic NP-EVs expressed HLA-G, as detected by ExoView and super-resolution microscopy. HLA-G, the distinct HLA isoform involved in maternal tolerance of the fetus, was previously reported to characterize EVs from the placenta *in vitro* or *in vivo* in the maternal circulation (Eunsung Mouradian, 2008; Orozco *et al*., 2009). It is also conceivable that the reported presence of HLA-G within the amniotic fluid could be related to its expression on the EV membrane (McMaster *et al*., 1998). The origin of the HLA-G expressing EVs in the amniotic fluid is unclear. They could possibly directly originate from the amniotic membrane surrounding the fetus and containing the amniotic fluid. Another possible explanation is the passage of trophoblast-derived EVs through the chorion and amnion to reach the amniotic fluid (McMaster *et al*., 1998). The placental origin of amniotic fluid-EVs is also suggested by the analyses of the deregulated miRNA target genes in preeclampsia, showing the most represented cell targets origin from the myoblast, the omentum and the placenta. The placental and stem cell origin of NP-EVs supports their use in regenerative medicine, as recently proposed (Mitrani *et al*., 2021).

Proteomic and miRNA profile analysis of amniotic fluid-EVs has been gaining an increasing interest in identifying disease-associated biomarkers and mechanisms (Dixon *et al*., 2018). In this study, we identified the specific increase in the levels of surface-associated CD105 in PE-EVs, whereas other surface markers showed similar expression levels. Endoglin (CD105) is a co-receptor of TGF-β1 and TGF-β3, highly expressed on the surface of syncytiotrophoblasts and endothelial cells (Valluru *et al*., 2011). An increase in soluble endoglin is a hallmark of preeclampsia (Mutter and Karumanchi, 2008; Schuster *et al*., 2020) and is responsible for endothelial damage by trapping soluble TGF-β molecules (Valluru *et al*., 2011). We here confirmed the increased CD105 levels through different orthogonal techniques. The use of a CD105-based capture chip allowed us to demonstrate that the increase in CD105 expression was not due to HLA-G expressing placental EVs. The origin of CD105 expressing EVs could be possibly attributed to cells of fetal or fetal membrane origin in the fluid, including AFSCs. Indeed, the analysis of cultured amniotic fluid mesenchymal stromal cells showed that the released EVs were largely CD105^+^, whereas the HLA-G^+^ EVs, representing a minor fraction, did not co-express CD105. It would be of interest in future studies to evaluate the CD105/HLA-G co-expression by cells isolated from preeclamptic amniotic fluid.

The increased CD105 level in PE-EVs is in agreement with previous data, showing higher levels of endoglin on EVs produced by placental explants treated with preeclamptic sera (Tannetta *et al*., 2013) and support the concept that surface receptors expressed by EVs may act as a decoy for soluble factors (Tannetta *et al*., 2013; Sedrakyan *et al*., 2017). In analogy, we found that PE-EVs affected angiogenesis *in vitro* and that this effect was abolished when CD105 was blocked by a specific antibody. A slight antiangiogenic effect was also observed for NP-EVs, in parallel with data showing that AFSC-EVs do not promote angiogenesis despite their regenerative effect (Takov *et al*., 2020). However, this was not prevented by the anti-CD105 antibody, suggesting the involvement of other factors. Previous data showed that administration of human EVs derived from preeclamptic placentas in mice resulted in damage to the vasculature, poor fetal nutrition and promoted endothelial permeability and glomeruli damage (Chang *et al*., 2018). In particular, the study provided an additional mechanism for the antiangiogenic effect of EVs, related to the presence of soluble endoglin transfer from the EV cargo to endothelial cells (Chang *et al*., 2018). Our results, showing the antiangiogenic effect of amniotic preeclamptic EVs, highlight a new role for EVs present in the amniotic fluid, possibly acting on both fetal and maternal compartments. Indeed, many reports described feto-placental endothelial dysfunction in preeclampsia, as a detriment to both the mother and the fetus (Escudero *et al*., 2016). It would be of interest to investigate the possible effect of PE-EVs on fetal development.

Several miRNAs with relevant roles in placenta development and angiogenesis have been detected in biological fluids and tissues of preeclamptic patients, including specific miRNA sets in the deriving EVs and are considered as a potential biomarker and “fingerprint” of placental dysfunction (Fallen *et al*., 2018; Gray *et al*.; 2017; Menon *et al*., 2019; Salomon *et al*., 2017; Ospina-Prieto *et al*., 2016; Escudero *et al*., 2016; Cronqvist *et al*., 2017). We here identified 46 miRNAs deregulated in amniotic fluid-derived PE-EVs, and we confirmed a differential expression in 6 miRNAs, involved in trophoblast proliferation and invasion (let-7 and miR-613), as well as in the modulation of placental function (miR-15, miR-26a) and angiogenesis (miR-195, and miR-144) (Cross *et al*., 2015, Hu *et al*., 2009, Bai *et al*., 2012 Guan *et al*., 2016 Choi *et al*., 2013 Li *et al*., 2013). Interestingly, although these miRNAs did not overlap with the EV miRNA profile alterations described in maternal circulation (Fallen *et al*., 2018; Gray *et al*.; 2017; Menon *et al*., 2019; Salomon *et al*., 2017), the majority (let-7, miR-15b, miR-195, and miR-26a-1) have been reported to be modulated in placental tissue of preeclamptic patients (Cross *et al*., 2015; Mayor-Lynn *et al*., 2011; Hu *et al*., 2009; Bai *et al*., 2012; Choi *et al*., 2013). This underlines a differential profile of amniotic fluid-EVs from those in the maternal circulation, and may further support their origin from placental deriving cells. These differentially expressed miRNAs may target genes involved in various functions, including development and angiogenesis, and are tightly involved in VEGF regulation and vascular remodeling. Indeed, deregulated pathways associated with angiogenesis, invasion and inflammation have been previously reported by target interaction analysis of miRNA content carried by preeclamptic maternal blood (Pillay *et al*., 2019, Salomon *et al*., 2017).

In conclusion, our study characterizes at single-EV level, EVs present in the amniotic fluid, an interesting biofluid with a possible relevance for therapeutic application. Amniotic fluid-derived EVs showed a heterogeneous origin, expressing markers of fetal and placental cells. We highlight the differential expression of antiangiogenic factors in normal and preeclamptic amniotic fluid-derived EVs, both at surface and cargo level. These characteristics may reflect the hypoxic and antiangiogenic general status of preeclampsia and could possibly play a role by affecting the developing fetus or the surrounding fetal membranes. Finally, EVs within the amniotic fluid could potentially be used as a diagnostic test if investigated at earlier stages of pregnancy.

## Acknowledgements

We thank Dr. Sharad Kholia for his help with the figure and manuscript editing and correction. We thank TRACON Pharmaceuticals (San Diego, CA) for providing the anti-CD105 Ab TRC-105.

