## Supplementary data for "Single extracellular vesicle analysis in human amniotic fluid shows evidence of phenotype alterations in preeclampsia"

***Supplementary Table 1.*** *Volume of amniotic fluid and the corresponding concentration of EVs as particles/ml of initial volume, based on nanoparticle tracking analysis.*

| | Normal pregnancies | | Preeclampsia | |
|---|---|---|---|---|
| Sample N. | Sample volume (ml) | Concentration (particles/ml) | Sample volume (ml) | Concentration (particles/ml) |
| **1** | 20 | - | 12 | 2.80E+09 |
| **2** | 10 | 4.5 E+09 | 46 | 3.48E+09 |
| **3** | 6 | 2.58E+09 | 11 | 1.13E+09 |
| **4** | 12 | 2.81E+09 | 15 | 2.21E+09 |
| **5** | 4 | 3.8E+09 | 15 | 1.03E+09 |
| **6** | 4 | 3.6E+09 | 15 | 5.02E+09 |
| **7** | 10 | 2.53E+09 | 15 | 3.30E+09 |
| **8** | 4 | 6.59E+08 | | |
| **9** | 19.5 | 2.75E+09 | | |
| **10** | 4.5 | 2.18E+09 | | |
| **11** | 22 | 3.18E+11 | | |
| **12** | 15 | - | | |
| **13** | 22 | 3.08E+09 | | |
| **14** | 21 | 2.61E+09 | | |
| **15** | 20 | 3.08E+09 | | |
| **16** | 10 | 4.5E+09 | | |
| **17** | 6 | 2.58E+09 | | |
| **18** | 12 | 2.81E+09 | | |
| **19** | 4 | 3.8E+09 | | |
| **20** | 4 | 3.6E+09 | | |
| **21** | 19 | 2.79E+09 | | |
| **22** | 15 | - | | |
| **23** | 19 | 3.65E+09 | | |
| **24** | 22 | 2.27E+09 | | |
| **25** | 10 | 4.5E+09 | | |
| **26** | 6 | - | | |
| **27** | 30 | - | | |
| **28** | 15 | 2.27E+09 | | |
| **29** | 19 | 4.9E+08 | | |
| **30** | 19 | 2.98E+09 | | |
| **31** | 15 | 2.27E+09 | | |
| **32** | 19 | 2.81E+09 | | |
| **33** | 22 | - | | |
| **34** | 10 | - | | |
| **35** | 25 | 6.776E+10 | | |
| **36** | 18 | 5.481E+10 | | |
| **37** | 22.5 | - | | |
| **38** | 41 | 3.914E+11 | | |
| **39** | 27.5 | - | | |
| **40** | 15 | - | | |

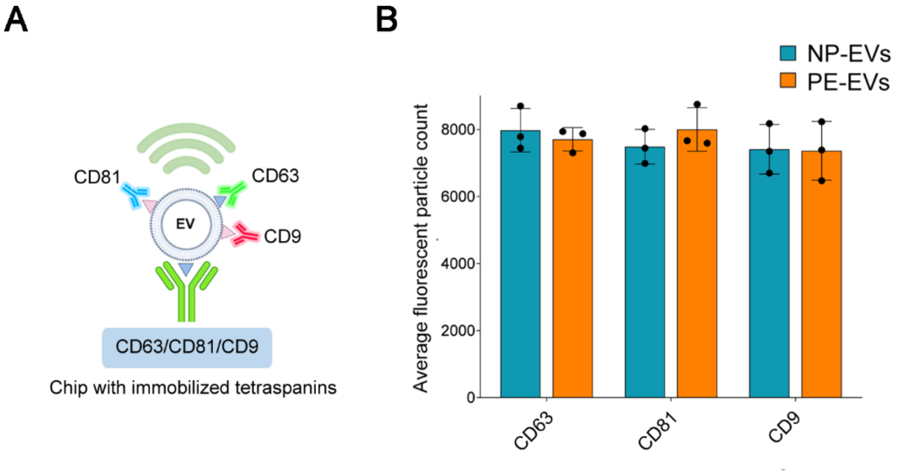


***Supplementary Figure 1*. ExoView analysis of amniotic fluid-derived NP-EVs (n=3) and PE-EVs (n=3).** *5.8x10$^{8}$ EVs in final volume of 35 µl of buffer were used for all samples.* ***A)*** *Image in panel B was created with https://biorender.com.* ***B)*** *ExoView chips spotted with single tetraspanins and detected with anti- CD63, CD81, CD9 fluorescent Abs confirmed a similar tetraspanins expression level between NP and PE-EVs. Three different samples were analysed for each experimental condition. No statistical differences were observed.*

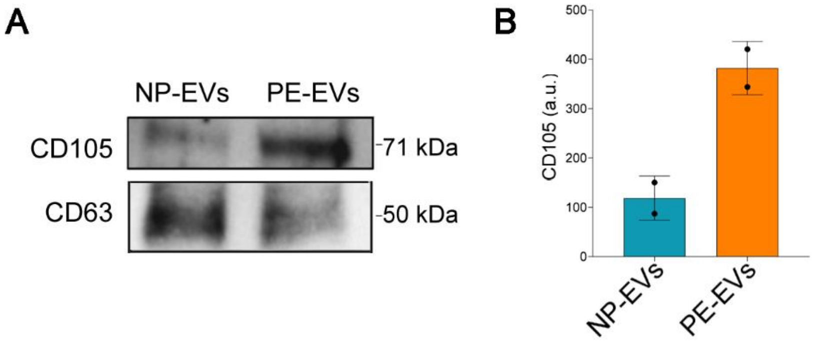


***Supplementary Figure 2****. Western Blot image and band intensity quantification showing CD105 expression differences between NP-EVs and PE-EVs.* ***A)*** *Representative image of the Western Blot showing the presence of CD105 and CD63 in NP- and PE-EVs.* ***B)*** *Quantification of Western Blot. CD63 was used as a loading control and the total lane intensity of CD105 was normalized to CD63. Three different NP-EV or PE-EV samples were pooled and used for the protein analysis.*

***Supplementary Table 2****. miRNA oligonucleotide sequences used for miRNA validation.*

| miRNA | Oligonucleotide sequence |
|---|---|
| hsa-miR-106a | AAAAGUGCUUACAGUGCAGGUAG |
| hsa-let-7e | CTATACGGCCTCCTAGCTTTCC |
| has-miR-141 | UAACACUGUCUGGUAAAGAUGG |
| hsa-miR-24 | UGGCUCAGUUCAGCAGGAACAG |
| hsa-miR-30d | CUUUCAGUCAGAUGUUUGCUGC |
| hsa-miR-29b | GCUGGUUUCAUAUGGUGGUUUAGA |
| hsa-miR-1247 | ACCCGTCCCGTTCGTCCCCGGA |
| hsa-miR-613 | AGGAATGTTCCTTCTTTGCC |
| hsa-miR-520c-3p | AAAGTGCTTCCTTTTAGAGGGT |
| hsa-miR-520b | AAAGTGCTTCCTTTTAGAGGG |
| hsa-miR-140-3p | TACCACAGGGTAGAACCACGG |
| hsa-miR-1290 | TGGATTTTTGGATCAGGGA |
| hsa-miR-195 | CCAATATTGGCTGTGCTGCTCC |
| hsa-miR-150 | TCTCCCAACCCTTGTACCAGTG |
| has-miR-144 | GGATATCATCATATACTGTAAG |
| has-miR-15b | UAGCAGCACAUCAUGGUUUACA |
| has-miR-19b | UGUGCAAAUCCAUGCAAAACUGA |
| has-miR-126 | UCGUACCGUGAGUAAUAAUGCG |
| has-miR-26a-1 | CCUAUUCUUGGUUACUUGCACG |
| has-miR-486 | CGGGGCAGCUCAGUACAGGAU |

***Supplementary Table 3****. Mean CT values from the miRNA array showing the top and bottom 10 expressed miRNAs in NP-EVs (n=3 NP-EV samples).*

| | Bottom 10 miRNAs | Mean CT | Top 10 miRNAs | Mean CT |
|---|---|---|---|---|
| **1** | hsa-miR-518 | 38.389 | hsa-miR-545 | 24.117 |
| **2** | hsa-miR-374 | 38.000 | hsa-miR-1274A | 27.890 |
| **3** | miR-93 | 38.000 | hsa-miR-1233 | 28.357 |
| **4** | hsa-miR-195 | 38.000 | hsa-miR-200c | 28.661 |
| **5** | hsa-miR-636 | 37.405 | hsa-miR-1260 | 28.919 |
| **6** | hsa-miR-190b | 37.338 | hsa-miR-574-3p | 29.099 |
| **7** | hsa-miR-151-3p | 37.157 | hsa-miR-1274B | 29.183 |
| **8** | hsa-miR-551a | 36.688 | hsa-miR-571 | 29.246 |
| **9** | hsa-let-7d | 36.364 | hsa-miR-720 | 30.041 |
| **10** | hsa-miR-32 | 36.192 | hsa-miR-191 | 30.340 |

***Supplementary Table 4****. Described role in pregnancy and preeclampsia (PE) of the 20 miRNAs chosen for validation.*

| | miRNA | Origin | Altered in PE | Possible function in PE | Ref. |
|---|---|---|---|---|---|
| **Downregulated** | **miR-15b** | Placenta | Yes | Regulation of placental renin−angiotensin associated genes. | Mayor-Lynn *et al.*, 2011 |
| | **miR-19b** | Plasma/plasma EVs | Yes | Placenta development | Kumar *et al.*, 2013, Sandrim *et al.*, 2016 |
| | **mir-26a-1** | Placenta | Yes | Placental inflammation | Choi *et al.*, 2013 |
| | **miR-30d** | Blood | - | Associated with hypertension. | Jia *et al.*, 2016 |
| | **miR-144** | Plasma | Yes | Regulates function in hypoxia. | Li *et al.*, 2013 |
| | **miR-150** | Umbilical cord derived EVs | In IUGR | Positive modulation of angiogenesis | Luo *et al.*, 2018 |
| | **miR-195** | Trophoblast cells | Yes | Downstream regulation of VEGF-A | Hu *et al.*, 2009, Bai *et al.*, 2012 |
| | **miR-486** | Plasma EVs | Yes | Suppression of placental development, angiogenesis and cell migration | Salomon *et al.*, 2017 |
| | **miR-520b** | Trophoblast cells | - | Regulation of placenta development | Takahashi, Ohkuchi and Usui, 2014 |
| | **miR-520c-3p** | Trophoblast EVs | - | Regulation of trophoblast invasion | Takahashi *et al.*, 2017 |
| | **miR-613** | Tumour cells | - | Suppression of proliferation and cell invasion | Guan *et al.*, 2016 |
| | **miR-1247** | Placenta | Yes | Regulation of placenta development | Enquobahrie *et al.*, 2011 |
| **Upregulated** | **miR-let-7e** | Trophoblast cells | - | Altered by oxidative stress in trophoblast cells. | Cross *et al.*, 2015 |
| | **miR-24** | Serum | - | cell growth, trophoblast differentiation, angiogenesis | Yang *et al.*, 2011 |
| | **miR-29b** | Trophoblast cells | - | increased apoptosis and decreased invasion and angiogenesis in trophoblast cells. | Cross *et al.*, 2015 |
| | **miR-106a** | Trophoblast cells | - | inhibition of trophoblast differentiation and syncytiotrophoblast formation. | Kumar *et al.*, 2013 |
| | **miR-126** | Endothelial cells | - | Modulation of angiogenesis | Guenther and Schrepfer, 2016 |
| | **miR-140-3p** | Tumor cells | - | Suppression of cell growth and increase of apoptosis | Jiang *et al.*, 2019 |
| | **miR-141** | Plasma Placenta Placental explant-derived EVs | Yes | Regulation of trophoblast invasion and angiogenesis | Ospina-Prieto *et al.*, 2016, Li *et al.*, 2013, Fallen *et al.*, 2018 |
| | **miR-1290** | Serum | Yes | Regulation of inflammation | Kim *et al.*, 2020 |

### Bioinformatic analysis

For the bioinformatic analysis of the deregulated miRNA present in PE-EVs, we first identified the validated targets, subsequently analysed the targets for functional enrichment analysis and target tissue distribution and finally we focused on the angiogenesis-related targets.

The search of gene targets identified more than 1000 strong evidence miRNA-gene interactions, involving 667 unique genes, being VEGF gene central in the interactome (Supplementary Figure 3).

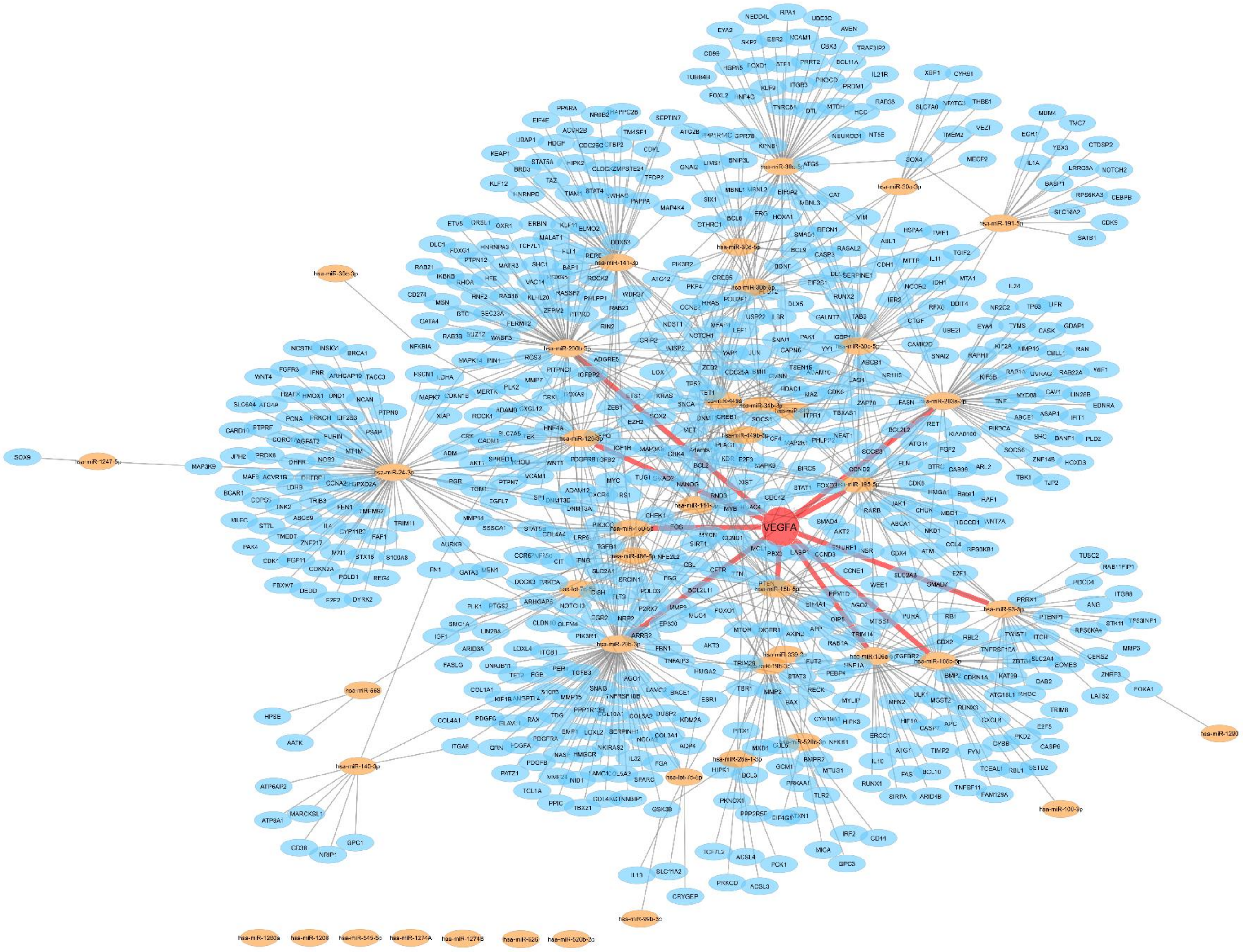


***Supplementary Figure 3.*** *Target interaction network miRNA (orange) - gene (blue), showing VEGFA as a central gene within the interactome (highlighted in red).*

Further functional enrichment analysis of GO terms and pathways supported the miRNA involvement in the regulation of a variety of pathways including development, angiogenesis and vascular remodelling. In particular, KEGG pathway enrichment analysis reveals 159 significantly enriched pathways. Most of the top identified pathways (P=<0.001) are cancer related. Enriched top non-cancer pathways are cellular senescence, PI3K-Akt signalling pathway, AGE-RAGe signalling pathway, and Focal adhesion. Reactome pathway enrichment analysis identified 14 significantly enriched pathways (P=< 0.001), most of which are general pathways, such as signal transduction, developmental biology, extracellular matrix organization, and immune system. More specific enriched pathways are cellular senescence, NGF and PDGF, SCF-KIT, and Fc epsilon receptor (FCERI) signalling. Wikipathways pathway enrichment analysis shows 296 significantly overrepresented pathways. The ten most significant pathways match the results of Reactome and KEGG results, with the addition of VEGFA-VEGFR2 signalling pathway (P=<0.001) (Supplementary Figure 4).

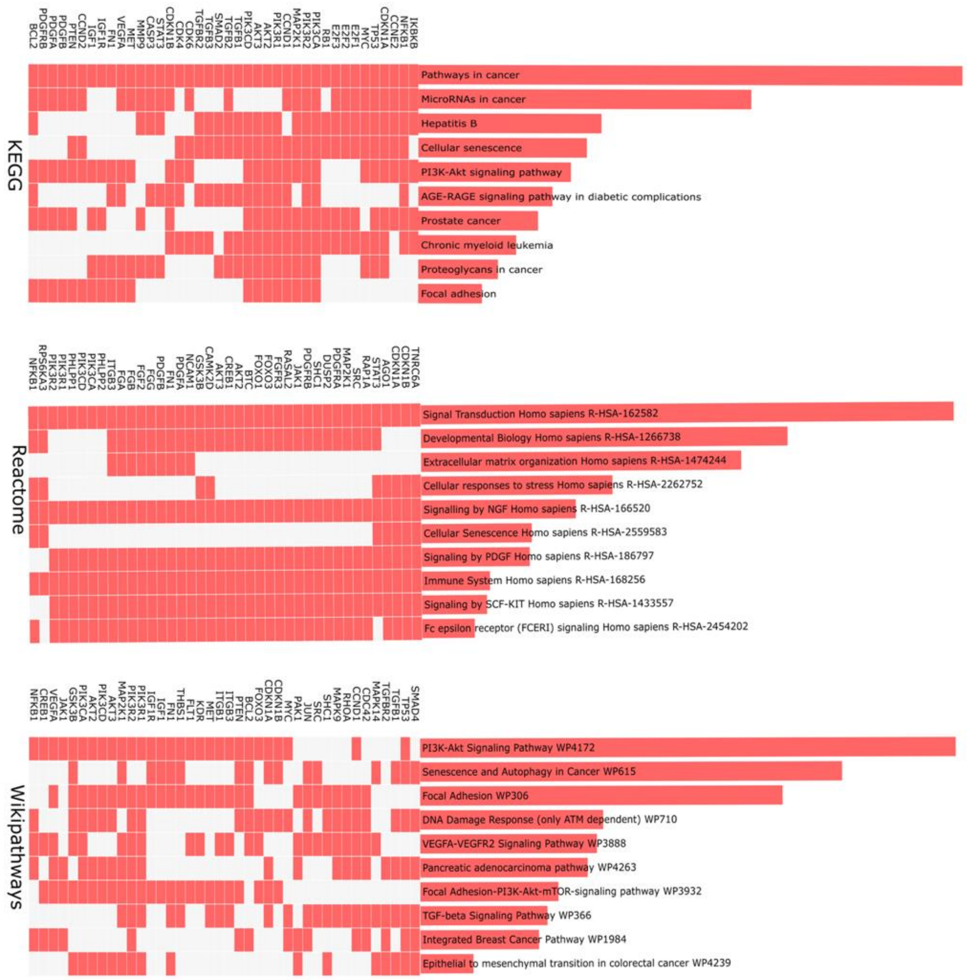


***Supplementary Figure 4:*** *Clustergrams of top 10 enriched pathways including the top 40 genes, falling into these pathways sorted by Padj in KEGG, Reactome, and Wikipathways. Pathway enrichment analysis was performed in the web interface EnrichR based on the databases KEGG Reactome, and Wikipathways (available under: https://maayanlab.cloud/Enrichr/enrich?dataset=c30948f97dc9dac1cc50f208f2503132). The*

*consideration of several databases was necessary, as they contain different representations of the same pathway (Mubeen et al., 2019).*

Accordingly, the analysis of target tissue and cell specific gene expression profile identified, among the top represented, myoblast, omentum and placenta (Supplementary Figure 5).

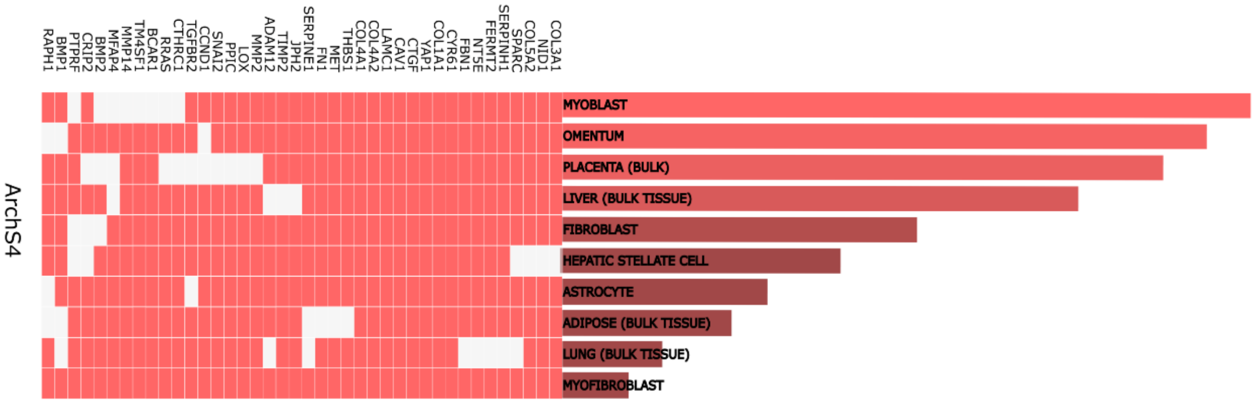


***Supplementary Figure 5.*** *Overview of Enrichment Analysis results sorted by database and -log10 (padj) ARCHS4 EnrichR tissue and cell targets.*

### Angiogenic profile of deregulated miRNAs in PE-EVs

We subsequently focused on the modulation by the PE deregulated miRNAs of angiogenic genes, by filtering for angiogenesis related terms the previous GO biological processes analysis. We show that many upregulated miRNAs targeted pro-angiogenic genes, suggesting an overall inhibition of the angiogenic process (Supplementary Figure 6A). Finally, we further analyzed the deregulated miRNAs for the identification of triplexes, that are miRNAs acting on a common target and resulting in an increased inhibitory effect, using TriplexRNA database. We identified six triplexes, with six unique targets (Supplementary Figure 6B). In particular, COL22A1 appeared targeted by cooperative miRNA with the lowest predicted free energy (-36.86), and energy gain (-15.08) of all identified cooperative miRNA complexes.

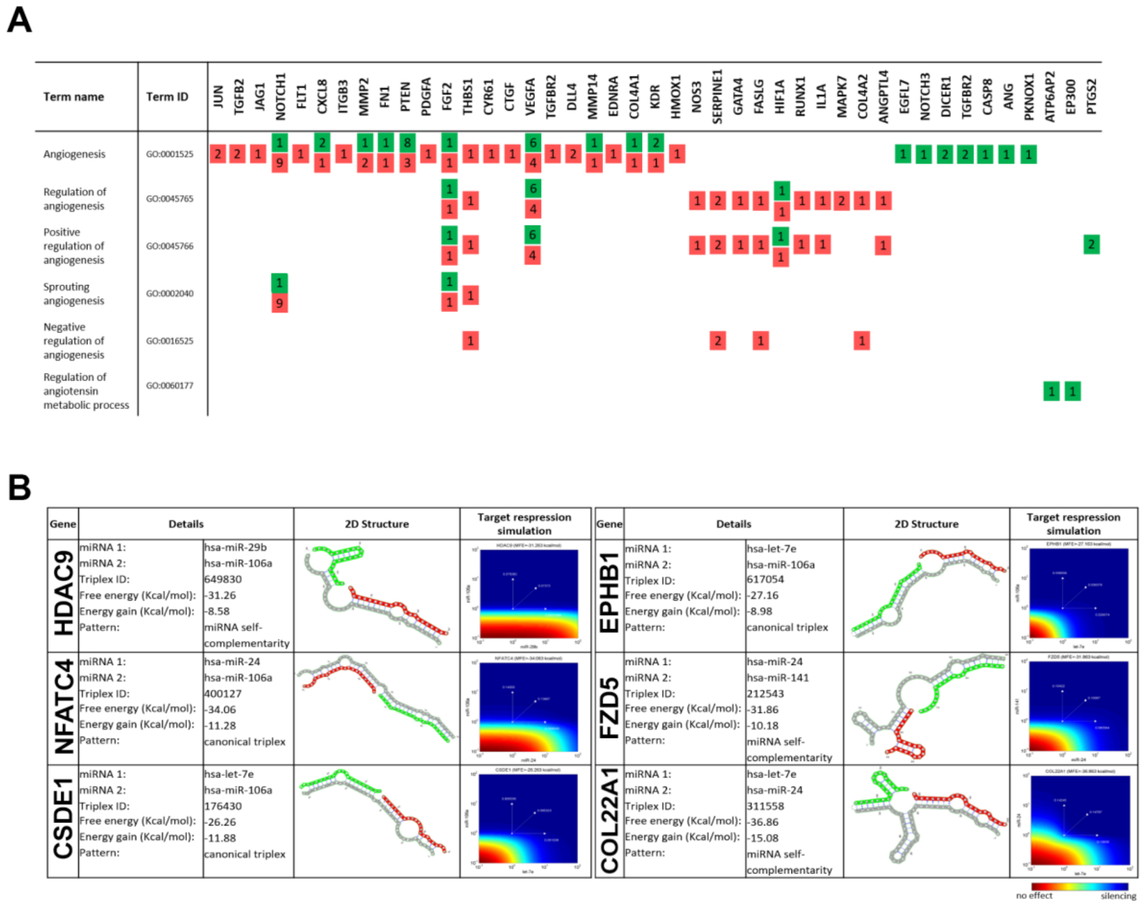


***Supplementary Figure 6. A)*** *Angiogenesis related GO terms and the genes targets of upregulated (red) and downregulated (green) miRNAs. The numbers on each square represent the number of miRNAs acting on each gene.* ***B)*** *Angiogenesis related RNA triplexes based on upregulated miRNAs. 2D structure is shown in the third column, with miRNA1 in red, miRNA2 in green, and target mRNA in grey. Target repression simulation can be seen in the fourth column where gene silencing by miRNA is visualized in blue and no effect in red.*

Several targets of upregulated miRNAs are of interest for further investigation, including NOX3, GATA4, RUNX1, IL1A, HMOX1, ANGPTL4, and the triplex target COL22A1. In particular, VEGF production appears regulated by NOX3, a reactive oxygen species generating NADPH oxidase (Carnesecchi *et al.*, 2006) GATA4, a stress-responsive regulator of angiogenesis, (Heineke *et al.*, 2007) as well as by RUNX1 (Whitmore *et al.*, 2021), JAK1 (Lin *et al.*, 2010), and IL1A (Fahey and Doyle, 2019) VEGFA further induces HMOX1 expression, leading to endothelial activation and angiogenesis (Bussolati and Mason, 2006). Finally, COL22A1 has been shown to maintain vascular stability in animal models (Ton *et al.*, 2018) and may be of interest to PE research as unresolved vascular remodeling has been implicated in PE (Schwartz, Vestweber and Simons, 2018). All these identified miRNAs might be acting in synergy on the angiogenic process (Supplementary Figure 7).

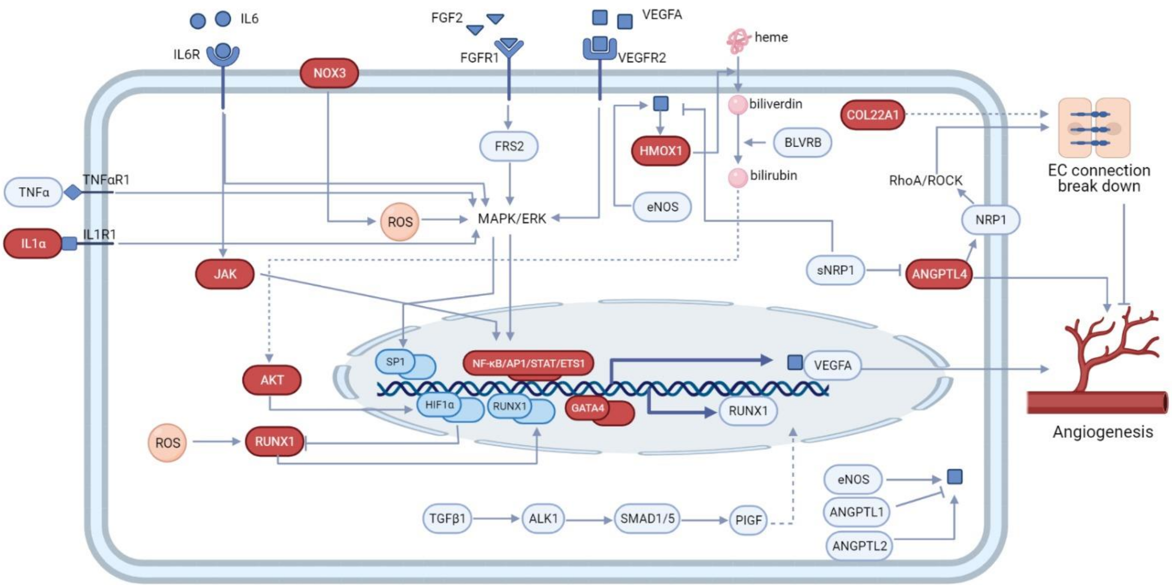


***Supplementary Figure 7:*** *Angiomodulatory signaling of significant gene target. Indicated gene targets are shown in red. Unknown interactions are indicated by dashed lines. For simplicity, gene instead of protein identifiers were used.*
